# AMPKα2 isoform mediates mTORC2 activation by glucose starvation in endothelial cells

**DOI:** 10.64898/2025.12.10.693448

**Authors:** Andrey N. Tikhonov, Nikita V. Podkuychenko, Vladimir P. Shirinsky, Asker Y. Khapchaev, Alexander V. Vorotnikov

## Abstract

AMP-activated protein kinase (AMPK) and the mechanistic target of rapamycin (mTOR) are central regulators of cellular metabolism. While AMPK is known to inhibit mTOR complex 1 (mTORC1), its role in regulating mTORC2 remains enigmatic, with recent evidence suggesting context-dependent activation. Here, we investigated the specific roles of AMPKα catalytic isoforms in coordinating mTOR signaling and functional metabolic adaptations in human umbilical vein endothelial cells (HUVECs) during glucose starvation. We found that loss of the major AMPKα1 isoform triggers an increase in otherwise minor AMPKα2 expression. While both isoforms are activated by glucose starvation and phosphorylate the canonical substrate acetyl-CoA carboxylase (ACC), only AMPKα2 is necessary and sufficient for transient activation of the mTORC2 signaling, as monitored by phosphorylation of downstream reporters, Akt and serum/glucocorticoid regulated kinase 1 (SGK1). This AMPKα2-dependent mTORC2 activation peaked at 30 minutes of starvation and then declined under prolonged starvation stress. Surprisingly, genetic ablation of the AMPKα2-mTORC2 axis through knockdown of AMPKα2, both AMPKα isoforms, or the essential mTORC2 component Rictor failed to attenuate the starvation-induced compensatory increase in glucose uptake. This adaptive response occurred robustly and identically across all genetic backgrounds. Collectively, our findings reveal an isoform-specific signaling module wherein AMPKα2 transiently activates mTORC2, but this pathway is functionally uncoupled from the critical adaptive response of glucose uptake, thus uncovering a remarkable resilience in the endothelial metabolic network and indicating that other, parallel pathways are the primary drivers of this essential survival mechanism.

## Introduction

Cellular metabolic homeostasis is paramount for organismal health, and its dysregulation underpins metabolic disorders such as type 2 diabetes and cardiovascular complications (Palmer & Clegg, 2022). The vascular endothelium, a critical interface between the blood and underlying У нас про Голодание Глюкозы, а здесь обе ссылки на избыток пальмитата и диабет, Где Глюкозы мноГо. Про Глюкозу бы что-нибудtissues, is highly susceptible to nutrient fluctuations and their excess, particularly saturated free fatty acids (Samsonov et al., 2022), resulting in endothelial dysfunction as a primary driver of diabetic complications (Goligorsky, 2017). In addition, hyperglycemia aggravates endothelial dysfunction (Shah & Brownlee, 2016), which makes glucose sensing important for endothelial function. Understanding molecular sensors that allow endothelial cells to adapt to the nutrient stress is therefore critical.

Two master regulators of cellular metabolism are AMP-activated protein kinase (AMPK) and mechanistic target of rapamycin (mTOR). AMPK is activated by phosphorylation sensitive to energy stress and AMP/ADP/ATP ratio or extracellular stimuli, and functions as an energy-sensing kinase to promote catabolic processes and inhibit the anabolic ones (Hardie, 2018). In contrast, mTOR operates within two distinct complexes, mTORC1 and mTORC2, and generally promotes anabolism and growth (Gonzalez et al., 2020). The AMPK and mTOR pathways are intricately linked, often in a reciprocal manner. A well-established paradigm is that AMPK phosphorylates and activates the inhibitory tuberous sclerosis complex 2 (TSC2) and/or phosphorylates and inhibits a scaffold mTORC1 subunit, Raptor, thereby suppressing the mTORC1 activity (Sarbassov et al., 2005).

The role of AMPK in regulating mTORC2 is less clear. As characterized by its core component Rictor, mTORC2 regulates cell survival, cytoskeletal organization, and metabolism primarily by phosphorylating the AGC kinases, Akt and serum/glucocorticoid regulated kinase 1 (SGK1), within their hydrophobic motif at Ser473 (Akt) and Ser422 (SGK1) (Sarbassov et al., 2005; García-Martínez & Alessi, 2008; Ragupathi et al., 2024). In SGK1, phosphorylated Ser422 primes further phosphorylation of Thr256 to activate the kinase (García-Martínez & Alessi, 2008). Recently, Kazyken et al. (2019) found that AMPK may directly phosphorylate mTORC2 to promote cell survival during acute energy stress, challenging the simple view of AMPK solely as an mTOR inhibitor.

AMPK exists as a heterotrimer with two catalytic α-subunits (α1 and α2). Although both isoforms are expressed in endothelial cells, the AMPKα1 is predominant and approximately 4-fold more abundant than AMPKα2 isoform (Colombo & Moncada, 2009; Khapchaev et al., 2024). The individual cellular functions of these isoforms are not fully delineated, including the endothelial cells. Furthermore, the specific AMPKα isoform responsible for engaging mTORC2 and physiological outcome of this interaction remain completely unknown.

In this study, we hypothesized that specific AMPKα isoforms differentially regulate mTORC1 and mTORC2 in endothelial cells to orchestrate an adaptive response to nutrient stress. We combined isoform-specific genetic knockdown and signaling analysis in HUVEC to dissect these pathways. We report that glucose starvation activates both AMPKα isoforms, but only AMPKα2 selectively activates the mTORC2-Akt/SGK1 axis. However, we find that within this pathway, AMPKα2-driven mTORC2 activation is functionally uncoupled from compensatory glucose uptake in response to glucose starvation, indicating that other mechanisms mediate this vital adaptive response to the nutrient deprivation.

## MATERIALS AND METHODS

### Cell Culture and Reagents

HUVECs were isolated from healthy donors, pooled, and cultured in M199 medium supplemented with ECGS on gelatin-coated dishes. Experiments were performed on passage 3-5 cells. Glucose starvation was induced by washing and incubating cells in DMEM containing 0-5 mM glucose for the indicated times. Control cells were incubated in DMEM with 5 mM glucose. All standard reagents, antibodies (against pACC-Ser79, acetyl-CoA carboxylase (ACC), pAkt-Ser473, Akt, pS6K1-Thr389, S6K1, pSGK1-Thr256, SGK1, AMPKα1, AMPKα2, Rictor, Raptor, mTOR, vinculin, glyceraldehyde-3-phosphate dehydrogenase (GAPDH), were obtained from Cell Signaling (USA).

### Lentiviral Transduction and shRNA Knockdown

Isoform-specific knockdown was achieved using lentiviral particles encoding shRNAs against AMPKα1, AMPKα2, both (pan-AMPKα), as previously described (Khapchaev et al., 2024). Silencing of Rictor was done in similar manner using 5’-GGTTTATTGACCACCCTTAGT as a target sequence. An empty vector not coding for any shRNA was used as a mock control. Cells were analyzed 5 days post-transduction to ensure efficient protein knockdown.

### Western Blotting

Cells were lysed in Laemmli buffer, and proteins were separated by SDS-PAGE, transferred to polyvinylidene fluoride (PVDF) membranes, and probed with specific primary and horse-reddish peroxidase-conjugated secondary antibodies. Chemiluminescence signals were developed using Clarity ECL kit (BioRad, USA) and detected using gel documentation system (Vilber Lourmat, France). ImageJ 1.54 freeware (NIH, USA) was used for densitometry to quantify images.

### Glucose Uptake Assay

Cells pre-incubated with oleate/palmitate were washed and placed in glucose-free DMEM medium for 10 min. Then, some cells were stimulated with 100 nM insulin for 15 minutes. Subsequently, 100 µM 2-deoxyglucose (2-DG) and 0.1 Ci/mol [3H]-2-deoxyglucose (USA, ARC) as a tracer were added to the cells for 10 minutes. The reaction was stopped by adding cold Hanks’ solution (PanEco, Russia) containing 4.5 g/l glucose. Immediately after this, cells were frozen. Cells were detached from the plates by adding RIPA lysis buffer. Samples were homogenized by passing through an insulin syringe. Samples were centrifuged at 13,000g for 10 minutes, the supernatant was transferred to scintillation vials with Ultima Gold scintillation fluid (PerkinElmer, USA), mixed, and the β-radiation of 2-DG was measured using a RackBeta counter (USA).

### Statistical Analysis

The data are presented as means ± SD of at least two independent experiments. The p values were calculated using Student’s *t* test or one-way ANOVA, as specified in the figure legends. Differences were considered statistically significant at p ≤ 0.05.

## RESULTS

### Glucose Starvation Activates AMPK and mTORC2 in Dose- and Time-Dependent Manner

To define an activation profile of the metabolic stress response of HUVEC to glucose starvation, we first analyzed the dose-dependence and time-course of AMPK and mTORC2 activation. We used phosphorylation of ACC at Ser79 (pACC-Ser79) and Akt at Ser473 (pAkt-Ser473) as direct markers of AMPK and mTORC2 activities, respectively (Fig. 1).

**Figure 1:**
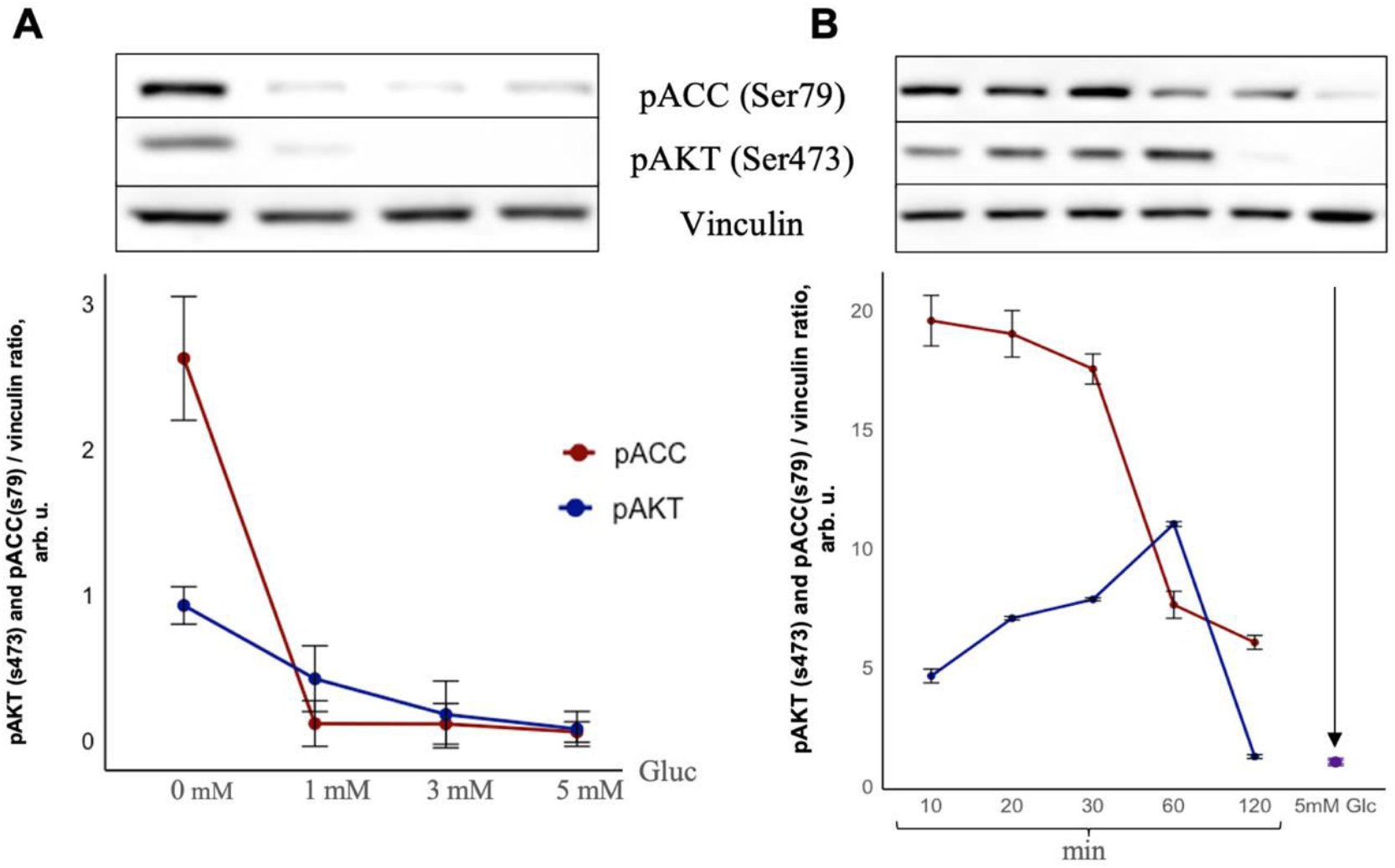
Glucose starvation activates AMPK and mTORC2 in a dose- and time-dependent manner. (A) Dose-dependent activation. HUVECs were incubated for 30 minutes in media titrated with the indicated concentrations of D-glucose (0, 1, 3, 5 mM). Cell lysates were immunoblotted for phosphorylated ACC (pACC-Ser79, red; AMPK activity) and phosphorylated Akt (pAkt-Ser473, blue; mTORC2 activity). Corresponding graphs (below) show densitometric quantification of the bands normalized to vinculin. (B) Time-dependent activation. HUVECs were subjected to full glucose starvation (0 mM) for the indicated times (10, 30, 60, 120 min). Lysates were immunoblotted and quantified for pACC-Ser79 (red) and pAkt-Ser473 (blue) as in (A). The data are presented as mean ± SD from at least 2 independent experiments.

First, we titrated the extracellular glucose concentration over a 30-minute period of cell treatment. Phosphorylation of ACC (pACC-Ser79) peaked at the lowest glucose level (0 mM) and progressively decreased as the glucose availability increased (Fig. 1A). This demonstrates that AMPK acts as a sensitive glucose sensor responding inversely to glucose concentration. Notably, phosphorylation of Akt (pAkt-Ser473), the canonical marker of mTORC2 activity, displayed a similar dose-response relationship. Its activation was strongest under severe starvation (0 mM Glc) and was completely abrogated at physiological glucose levels (5 mM) (Fig. 1B). This parallel dose-dependence provided indication that AMPK and mTORC2 activities are co-regulated by glucose availability.

We next performed a time-course experiment under conditions of full glucose starvation (0 mM) to resolve the kinetic relationship between these two pathways. AMPK activation was rapid and sustained, with a significant increase in pACC-Ser79 detectable within 10 minutes of starvation and slowly decreased for the full 120-minute duration (Fig. 1B). In contrast, mTORC2 activation followed a delayed kinetic profile. That is, pAkt-Ser473 levels began to rise from first 10 minutes of starvation and continued to increase, reaching a peak in 60 minutes marker (Fig. 1B).

Together, these data reveal a coordinated response: glucose starvation dose-dependently co-activates AMPK and mTORC2, with AMPK activation preceding and potentially upstream of the delayed mTORC2 response. Based on these kinetic and dose-response profiles, we selected 0 mM glucose and a 30-minute time point for subsequent experiments to ensure robust and specific activation of both AMPK and the initial phase of mTORC2 signaling.

### Lentiviral shRNA-Mediated Knockdown Reveals Reciprocal Regulation of AMPKα2 Isoform

Our study was designed to interrogate the specific functions of AMPKα isoforms during short-term metabolic stress. As a robust and effective gene silencing approach, we selected a lentiviral shRNA-mediated knockdown to achieve permanent gene silencing (Stewart, S.A., et al. 2003).

Using this system, we selectively knocked down AMPKα1, AMPKα2, or both isoforms (pan-AMPKα). Fig. 2 confirms the efficacy of this approach. The shRNA to AMPKα1 (shAMPKα1) completely reduced the protein signal, whereas it was evident in the mock-transduced and shRNAα2 or shRNAα1/2 cells. A striking and consistent observation was that the loss of AMPKα1 triggered a compensatory several-fold increase in the protein level of AMPKα2 isoform (Fig. 2A, and Fig. 2C). Consequently, the total AMPKα signal in these cells, as detected by pan-AMPKα antibody, predominantly represented the upregulated AMPKα2, although it was decreased as compared to that in mock cells or cells with knockdown of AMPKα2 (shAMPKα2). In contrast, the latter cells demonstrated unaltered levels of AMPKα1, possibly because of too little change due to a high initial content of AMPKα1 over low abundance of AMPKα2 isoform. As expected, shRNA targeting both isoforms (shAMPKα1/2) successfully reduced the levels of both AMPKα1 and AMPKα2, resulting in a near-total ablation of AMPK expression, as detected by pan-AMPKα antibody (Fig. 2A), and its signaling capacity (Fig. 3 below). This successful generation of isoform-specific AMPKα deficient HUVEC provided the essential genetic toolkit to dissect the unique roles of AMPKα isoforms in regulating mTOR signaling in subsequent experiments.

**Figure 2:**
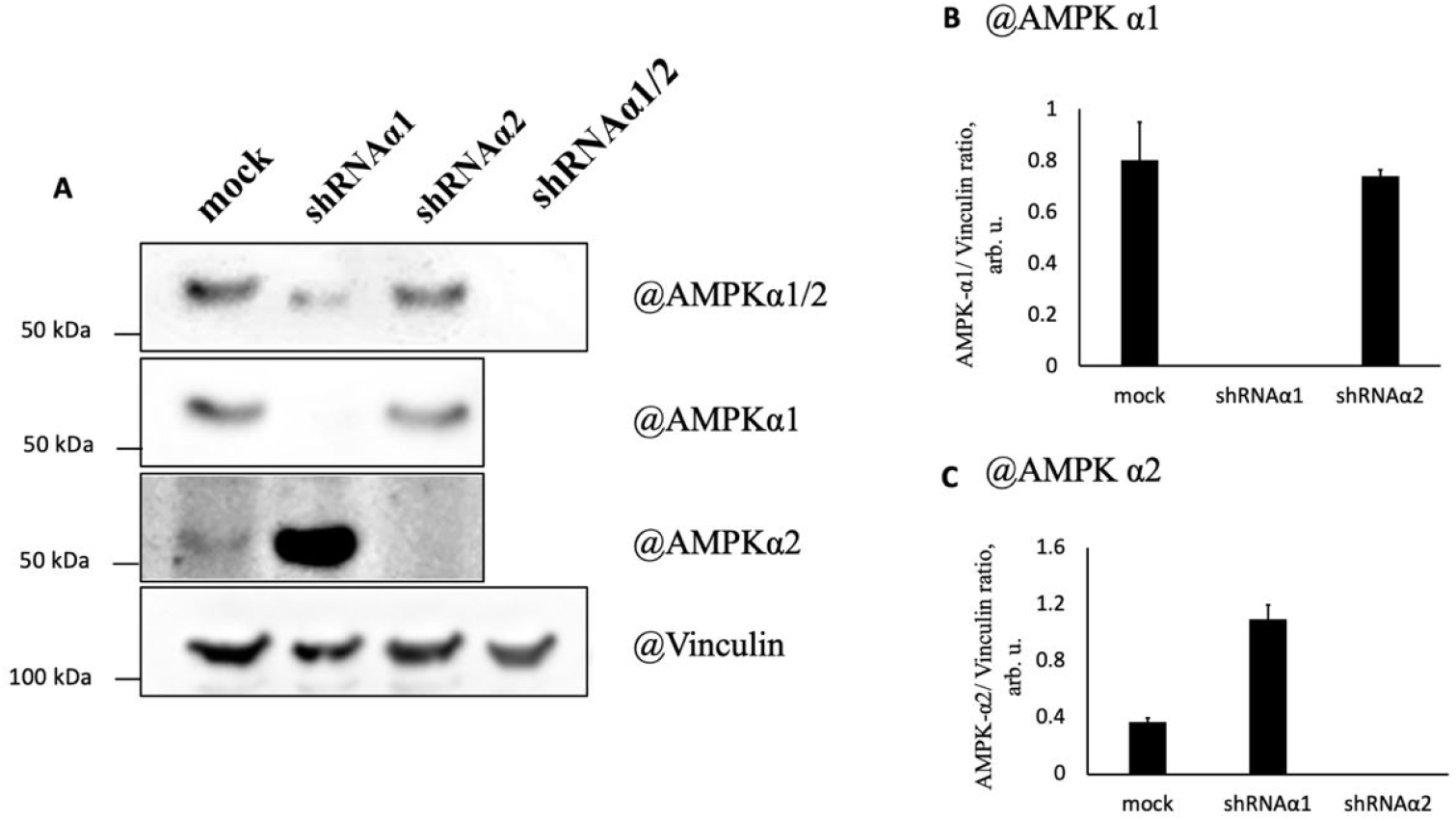
Lentiviral shRNA-mediated knockdown of AMPKα isoforms reveals compensatory upregulation of AMPKα2. (A) Representative Western blots of cell lysates after HUVEC transduction with empty lentivirus (Mock) or shRNAs to individual or both AMPKα isoforms as indicated on the top. The western membranes were probed for individual AMPKα isoforms, or total AMPKα (pan) as indicated on the right. Vinculin served as a protein loading control. (B-C) Densitometric quantification of AMPKα1 (B) and AMPKα2 (C) protein levels normalized to the vinculin content from (A). Data are presented as mean ± SD, n=4.

**Figure 3:**
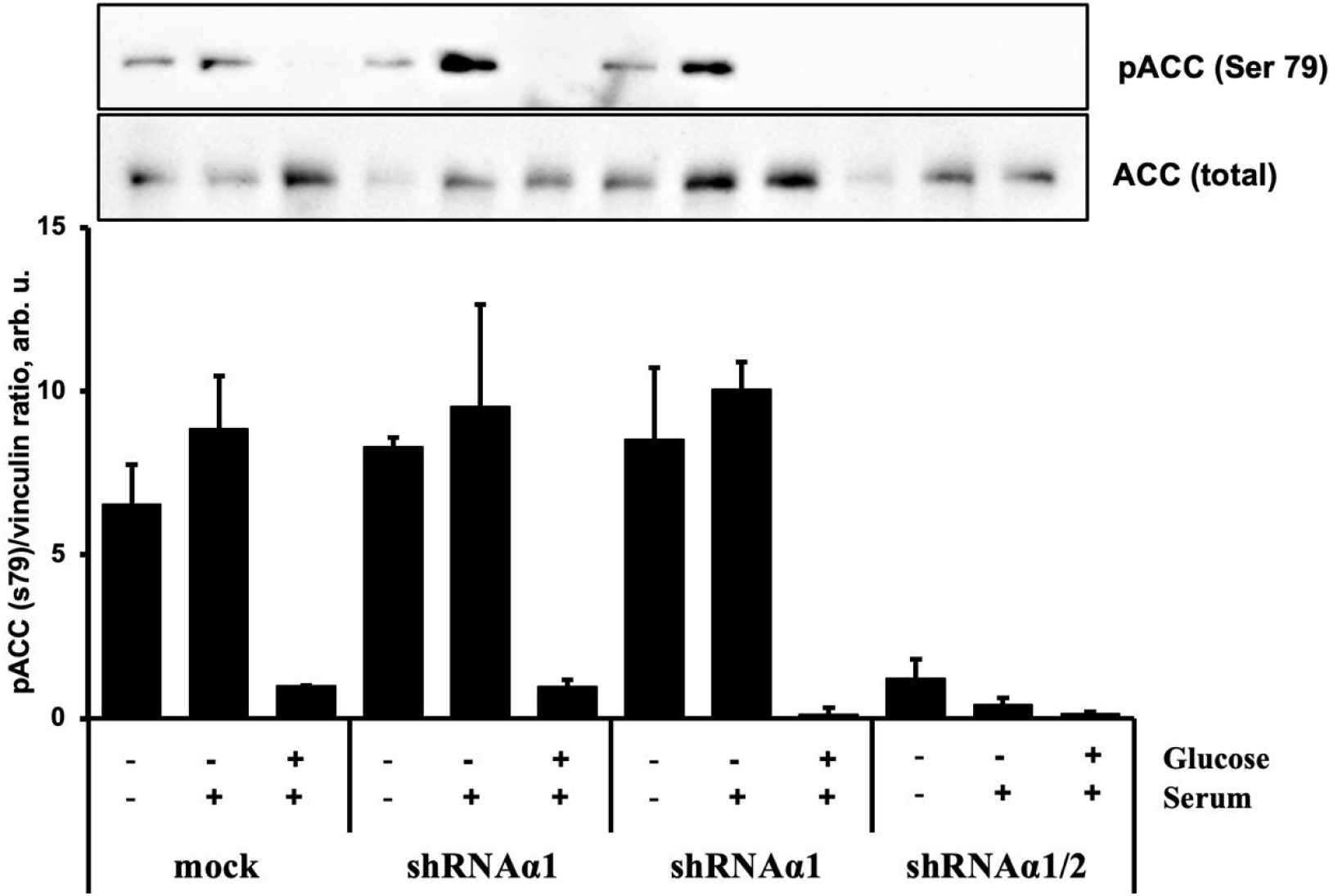
Glucose starvation activates both AMPKα isoforms independently of growth factors. Control (Mock) and AMPKα silenced HUVEC were incubated for 3 hours in the media containing either 5 mM or 0 mM glucose, in the presence or absence of serum/growth factors (FBS/ECGS) as indicated in the legend underneath. Shown are representative Western membranes probed for phosphorylated ACC (pACC-Ser79) (top) and total ACC (bottom) along with densitometric quantification of the pACC-Ser79/ACC ratio normalized to vinculin derived from 4 independent experiments

### Both AMPKα Isoforms Are Activated by Glucose Starvation and Phosphorylate ACC

We confirmed that our starvation protocol effectively activated either of AMPKα isoforms. Consistent with our initial characterization, glucose deprivation robustly increased phosphorylation ACC at Ser79 across all cell lines that retained at least one AMPKα isoform, the effect even lasting for 3 hours after glucose withdrawal (Fig. 3). Crucially, AMPK activation occurred to a similar extent regardless of the presence or absence of serum and growth factors (FBS+ECGS), underscoring that glucose availability, but not that of growth signals, is the primary trigger for AMPK activation under acute glucose starvation (Fig. 3).

As expected, the starvation-induced phosphorylation of ACC was completely abolished in pan-AMPKα knockdown cells (shАМРКα1/2), which lack both catalytic isoforms. This result confirms specificity of Ser79 in ACC as a reporter of total AMPK activity and, importantly, demonstrates that both AMPKα1 and AMPKα2 are functionally competent and can be activated by energy stress in endothelial cells.

### mTORC1 Activity Depends on Growth Factors but is not Acutely Regulated by AMPK in HUVEC

Consistent with its role as a sensor of growth signal, the mTORC1 activity assessed by phosphorylation of Thr389 in S6K1 strongly depended on the presence of fortified serum (FBS+ECGS), but was largely unaffected by glucose availability alone over a 3-hour period (Fig. 4). Furthermore, the activity of mTORC1 was not significantly altered by knockdown of any AMPKα isoform, indicating that AMPK does not exert a strong inhibitory effect on mTORC1 under these acute starvation conditions, regardless of knocking down any of its catalytic subunit isoform.

**Figure 4:**
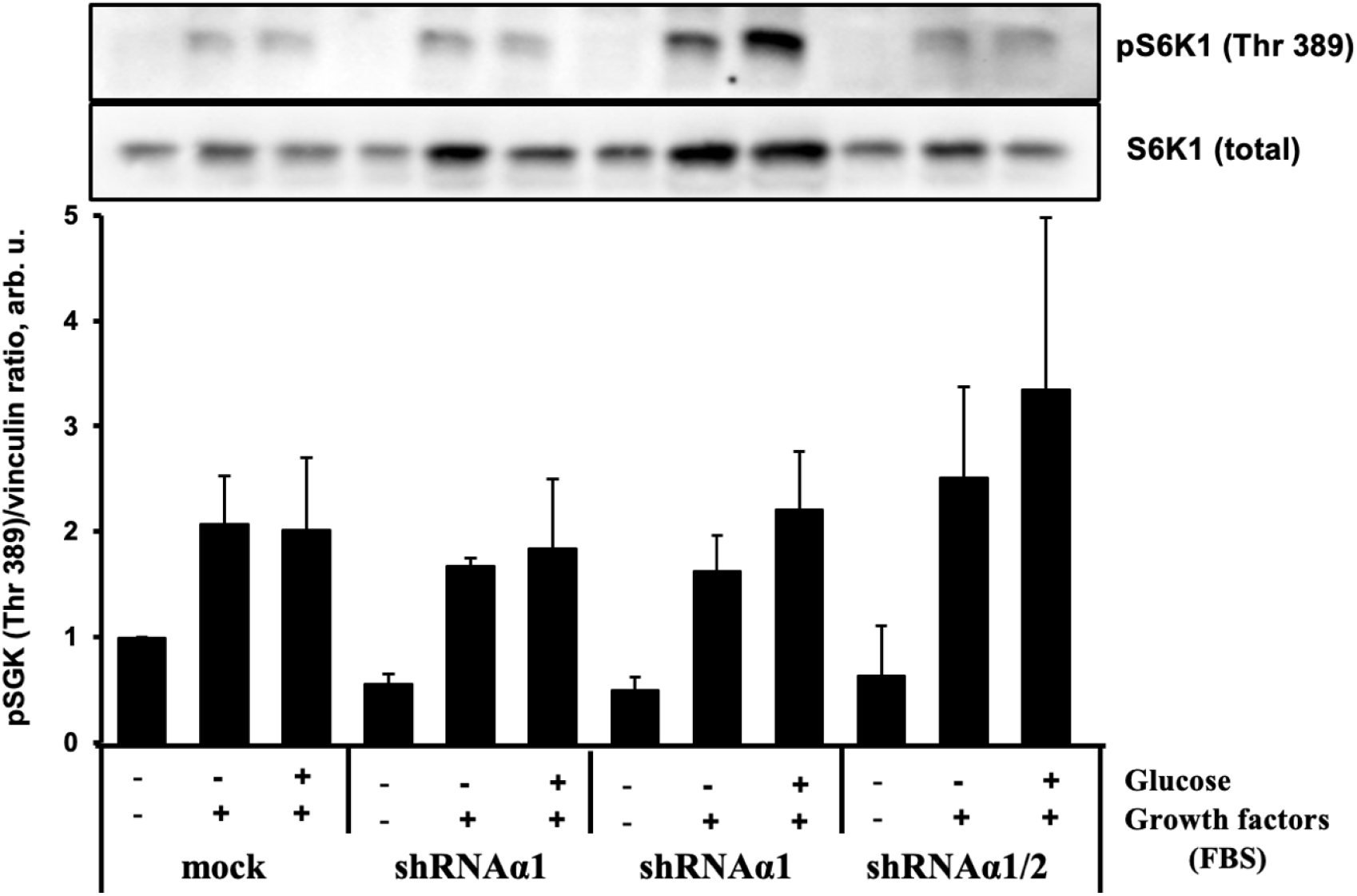
mTORC1 activity is dependent on growth factors but independent of AMPKα isoforms. Control (Mock) and AMPKα silenced HUVEC were incubated for 3 hours under the indicated conditions of glucose (5 mM or 0 mM) and serum fortified by endothelial growth factors (FBS+ECGS). Shown are representative Western filters probed for phosphorylated S6K1 (pS6K1-Thr389) and total S6K1 along with densitometric quantification of the pS6K1/S6K1 ratio. Data are presented as mean ± SD from 3 independent experiments.

### The AMPKα2 Isoform Selectively Activates mTORC2 upon Glucose Starvation

Having established that AMPK and mTORC2 activities were nearly maximal at 30 minutes of glucose starvation, as judged by ACC and Akt phosphorylation (Fig. 1B), we sought to definitively link this response to specific AMPKα isoforms. We therefore analyzed phosphorylation of Akt-Ser473 in our panel of AMPKα-deficient HUVECs after a 30-minute glucose starvation.

This analysis revealed that a potent and specific increase in Akt phosphorylation was exclusively contingent on the AMPKα2 isoform (Fig. 5A). Most strikingly, this activation was dramatically enhanced only in shAMPKα1 cells, which overexpress AMPKα2. Conversely, the starvation-induced response was completely abolished in cells lacking AMPKα2 (shAMPKα2 and shAMPKα1/2 cells). Taken together, this demonstrates that the magnitude of early mTORC2 activation is most likely a function of AMPKα2 abundance.

**Figure 5:**
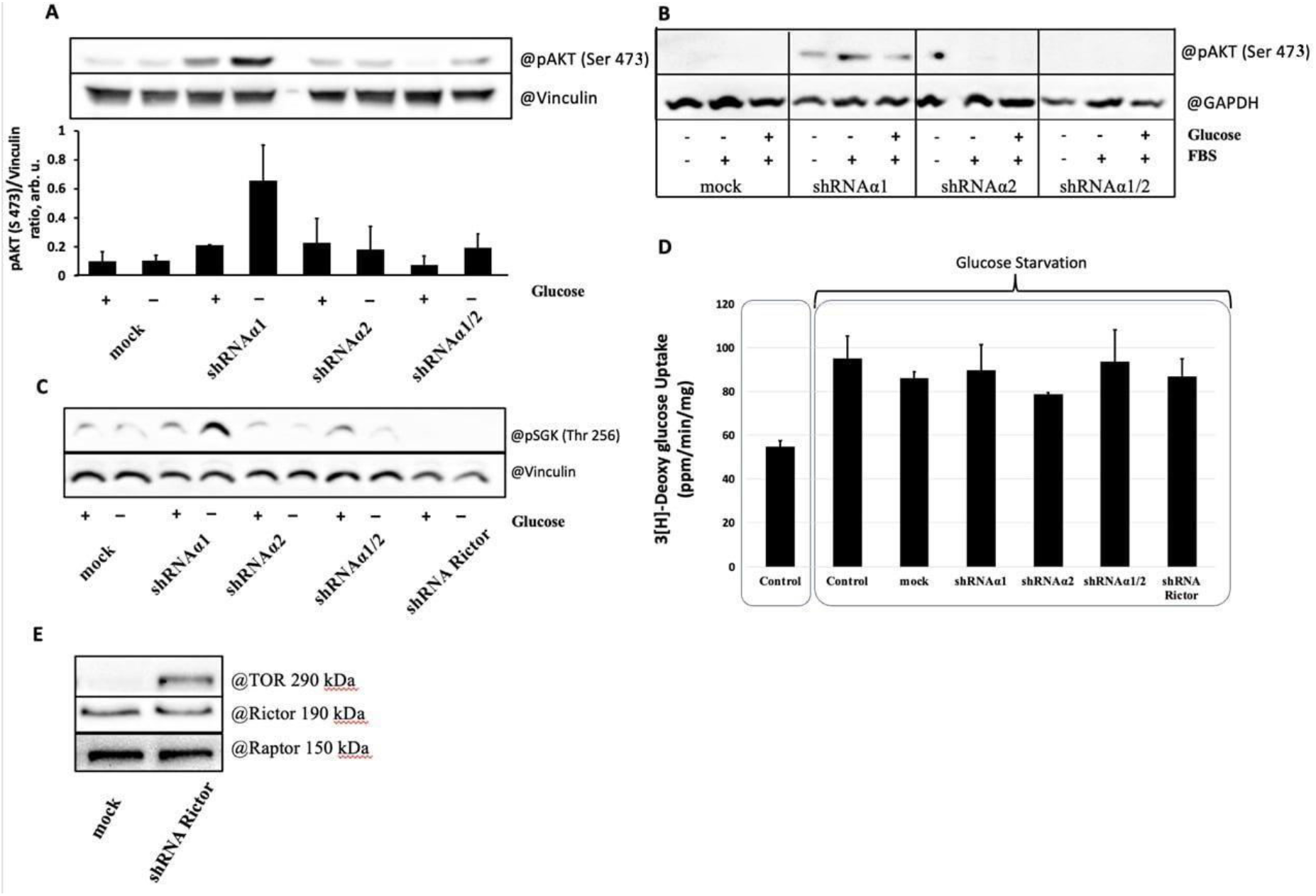
AMPKα2 is necessary for increased Akt phosphorylation upon glucose starvation. HUVEC were transduced by lentiviruses coding for either shAMPKα1, shAMPKα2, shAMPKα1/2, shRictor or mock control as indicated and subjected to deprivation of glucose for 30 min (A, C) or 3 hours (B, D). Cell lysates were analyzed by Western for molecular targets as indicated. (A) Ser473 phosphorylation in Akt after 30-minute glucose starvation. (B) Ser473 phosphorylation in Akt after 3-hour combined glucose and serum/growth factor starvation. (C) pSGK1-Thr256 phosphorylation after 3-hour glucose starvation. (D) Glucose uptake, measured by [^3^H]-2-deoxyglucose (2-DG) incorporation, in response to 30-minute starvation across all HUVEC variants, including Mock, shAMPKα1, shAMPKα2, shAMPKα1/2 (pan), and shRictor. Data are mean ± SD; *p < 0.05 vs. respective fed control; N.S. (not significant) for all comparisons between starved groups.

We next investigated the durability of this AMPKα2-dependent signaling. We hypothesised that the increased amount of AMPKα2 in HUVEC may enhance activation of mTORC2 and prolong its signaling to Akt. Thus, we extended the starvation period to 3 hours and assessed the impact of concurrent growth factor deprivation. Under these prolonged stress conditions, the AMPKα2-specific pattern of pAkt-Ser473 was again evident, although the starvation effect was much less apparent, possibly due to subsided mTORC2 activation and its uncoupling from stimulation by growth factors from serum (Fig. 5C). To confirm that this isoform-specific regulation targeted mTORC2 itself and not just Akt, we probed for phosphorylation of another mTORC2 substrate, SGK1, after 3 hours of glucose starvation. We assessed phosphorylation of Thr256 in the activation loop of SGK1, which is strongly dependent on phosphorylation of Ser422 in the hydrophobic motif by mTORC2 (García-Martínez & Alessi, 2008). Consistent with the Akt data, the pSGK1-Thr256 signal was specifically increased by glucose starvation in cells expressing high levels of AMPKα2 and was completely absent both in resting and stimulated mTORC2-deficient cells (Fig. 5C). mTORC2 deficiency was attained by shRNA against Rictor, a key scaffold protein for the mTORC2 assembly. Fig. 5E shows that Rictor was knocked down to undetectable levels while neither Raptor, the mTORC1 scaffold, nor the mTOR kinase expression were not altered. Altogether, these results are consistent with AMPKα2-driven activation of mTORC2 and further demonstrate that availability of AMPKα2 may impact the duration of the response to acute glucose starvation, which normally lasts for about an hour (Fig. 1B).

Collectively, these data provide conclusive evidence that AMPKα2 is the necessary and specific isoform responsible for the initial, transient activation of mTORC2 in response to acute glucose starvation. However, this specific signaling module appears to be a short-lived response that diminishes under prolonged stress.

### The AMPKα2-mTORC2 Axis is Uncoupled from Glucose Uptake Increased by glucose starvation

Given the clear role of AMPKα2 in initiating mTORC2 signaling, we sought to determine the functional consequence of this pathway for endothelial cell adaptation. We hypothesized that the AMPKα2-mTORC2 axis would be critical for the increase in glucose uptake, a potential key survival response to the nutrient deprivation. To this end, we measured the glucose uptake by control and starved HUVEC across our panel of knockdown cells using [^3^H]-2-deoxyglucose (2-DG).

As expected, 30 minutes of glucose starvation triggered a significant ~1.8-fold increase in 2-DG uptake in control cells (Fig. 5D), which confirmed that endothelial cells exert a rapid adaptive response to glucose starvation. However, this increase in glucose uptake was completely unaffected by the loss of any specific AMPKα isoform or by their combined knockdown (Fig. 5D). Furthermore, ablation of the essential mTORC2 component Rictor also failed to attenuate the starvation-induced glucose uptake. The magnitude of the increase was identical across all genetic backgrounds. These data demonstrate that while the AMPKα2-mTORC2 axis is activated by acute glucose starvation, it is not the primary pathway responsible for mediating the increase in glucose uptake and other pathways mediate this vital adaptive response independently of AMPKα2 and mTORC2.

## Discussion

Our study provides a dissection of AMPK isoform-specific signaling in endothelial cells under metabolic stress associated with lack of glucose availability. The main finding of our study is delineating a novel AMPKα2-mTORC2 axis. We revealed this signaling pathway in HUVEC using the AMPK catalytic isoform-specific knockdown and tracking the mTORC2 activity by phosphorylation of its downstream targets, Akt and GSK1. Our findings add a layer of specificity to the AMPK-dependent mTORC2 activation mechanism, which was earlier shown to contribute to increased cell survival under acute metabolic stress (Kazyken et al., 2019). Thus, results of our study are in agreement with the report by Kazyken et al. (2019) confirming the validity of the AMPK-mTORC2 axis in endothelial cells and extending the paradigm further to identify AMPKα2 as a responsible isoform. Here, we used a discriminating approach targeting individually either AMPKα1 or AMPKα2, versus simultaneous knockdown of both isoforms. This approach allowed us to find that AMPKα2 is the isoform uniquely responsible for transient activation of the mTORC2 pathway upon acute glucose starvation (Fig. 5A-C). Accumulating evidence indicates that the two AMPKα isoforms display both common and distinct functions (Yang et al., 2018; Yang et al., 2022; Rana et al., 2020). In our study, a widely acknowledged specific marker of AMPK activity, phosphorylation of ACC Ser-79, exemplifies a function common to both AMPKα isoforms (Fig. 3) while it contrasts with the AMPKα2-specific mTORC2 signaling activation. The study by Kazyken et al. (2019) used non-endothelial cells and reported that both AMPKα isoforms may activate mTORC2 based on simultaneous downregulation or ablation of both AMPKα isoforms. In other experimental settings, activation of AMPK was even associated with repressed mTORC2 activity in Th2 cells (Pandit et al., 2022) and myeloma cells (Wang et al., 2018). Therefore, the variability of the responses may be both cell type- and context-dependent, as we and Kazyken et al. (2019) observed the positive AMPK-mTORC2 axis under conditions of energetic stress.

Notably, in our experiments, effective downregulation of AMPKα1 expression in HUVEC was associated with a several-fold increase in the AMPKα2 expression (Fig. 2). This is in compliance with our previous findings (Khapchaev et al., 2024) and reports from other groups (Bess et al., 2011; Elsaie et al., 2021). Thus, in our model, the level of AMPKα2 expression was significantly higher than in standard HUVEC (Fig. 2) and may have contributed to strengthen the AMPKα2-mTORC2 axis. Though AMPKα2 is a minor isoform in most endothelial cells and, in particular, in HUVEC (Colombo & Moncada, 2009; Khapchaev et al., 2024), pulmonary artery endothelial cells represent an exception and express comparable levels of both AMPKα isoforms (Teng et al., 2013). In skeletal and cardiac muscle, AMPKα2 is a predominant isoform (Viollet et al., 2009) and in these cells, the role of the AMPKα2-mTORC2 axis may be physiologically justified to promote cell survival during the periods of an urgent need in high energy consumption on the background of nutrient shortage.

As noted above, the AMPKα2-mTORC2 axis was evident under conditions of acute deprivation of glucose (Fig. 5). Depriving cells of glucose is a well-known stress factor to activate AMPK (Salt et al., 1998). Glucose deprivation promotes translocation of AMPK from the cytoplasm to the lysosome and the mechanism operates with AMPK complexes containing either α-subunits (Zhang et al., 2017). On lysosomes, AMPK phosphorylates Raptor, a key scaffold component of mTORC1, and may thereby prevent inactive mTORC1 interaction with Raptor and mTORC1 activation (reviewed in Lin & Hardie, 2018). Endothelial cells also possess this mechanism with both AMPKα isoforms acting as mTORC1 repressors (Doshi et al., 2023). In our setting, the mTORC1 activity remained largely unaltered (Fig. 4). This may be due to acute AMPK activation, which was not long enough to activate the lysosome-mediated mTORC1 targeting.

In contrast, the exact mechanism how AMPK activation may converge to either activation or inactivation of mTORC2 is currently unclear. Direct phosphorylation of mTOR at Ser1261 was demonstrated in skeletal muscle, but had no effect on the mTORC2 activity (Li et al., 2025). Probably, a component of mTORC2 may be involved as it is the case with mTORC1 component Raptor. Identification of an AMPKα2-specific molecular target, which contributes to mTORC2 activation, requires further experimental research.

Glucose deprivation induced relatively rapid AMPK activation, and both AMPKα isoforms were simultaneously and comparably activated (Fig. 1, Fig. 3). As a functional response to acute glucose deprivation, we utilized the glucose uptake test. Deprivation of glucose is known to dramatically increase glucose transport and mTORC2 mediates insulin-stimulated glucose uptake in metabolic tissues (Kumar et al., 2004; Albert et al., 2016). Despite endothelium is not a conventional metabolic tissue, in HUVEC, we also observed an increase in glucose uptake upon deprivation of glucose (Fig. 5D). Studies on the AMPK-dependent glucose uptake in endothelial cells are limited and yielded conflicting results. That is, in endothelial cells, AICAR-dependent AMPK activation decreases the glucose uptake (Dagher et al., 1999; Dagher et al., 2001), while the shear stress-mediated AMPK activation increases the glucose uptake (Cronin et al., 2024). The most significant and unexpected finding of our work, however, is that the specific AMPKα2-mTORC2 signaling module is dispensable for the increase in glucose uptake. While we observed a robust and reproducible transient activation of mTORC2 by AMPKα2 at the 30-minute time point, genetic ablation of this entire axis—through knockdown of AMPKα2, all AMPKα, or Rictor—had no effect on the cell’s ability to enhance glucose transport in response to starvation. This result clearly uncouples the observed kinase cascade from the functional metabolic output we initially hypothesized it controlled.

This discrepancy suggests several intriguing possibilities. The AMPKα2-mTORC2 axis may regulate alternative adaptive processes crucial for survival under nutrient stress, such as cytoskeletal remodeling, autophagy, or the regulation of other metabolic enzymes, which were not in the focus of this study. The transient nature of the activation supports a role in initiating a specific, early adaptive program rather than a sustained chronic response to restore the nutrient transport. It is likely that redundant pathways ensure the critical function of glucose uptake.

Our data also reinforce the sophisticated segregation of metabolic signaling. We show that mTORC1 remains responsive to growth factors but is insulated from acute glucose fluctuations, consistent with its role as a growth promoter. Meanwhile, mTORC2 demonstrates sensitivity to nutrient status, albeit transiently, through AMPKα2. The functional independence of glucose uptake from this pathway indicates that endothelial cells are equipped with a resilient, multi-layered network to maintain energy homeostasis, where the failure of one signaling arm can be compensated by others.

## Acknowledgements

This work was supported by RSF grant #23-75-00027 (https://rscf.ru/project/23-75-00027/).

